# Highly differentiated CD4 T cells Unequivocally Identify Primary Resistance and Risk of Hyperprogression to PD-L1/PD-1 Immune Checkpoint Blockade in Lung Cancer

**DOI:** 10.1101/320176

**Authors:** Miren Zuazo-Ibarra, Hugo Arasanz, Gonzalo Fernández-Hinojal, Gato-Cañas María, Berta Hernández-Marín, Maite Martínez-Aguillo, Maria Jose Lecumberri, Angela Fernández, Lucía Teijeira, Ruth Vera, Grazyna Kochan, David Escors

**Author notes:** Correspondence to: Dr David Escors, or; Dr Grazyna Kochan,; Dr Ruth Vera. These authors contributed equally (Miren Zuazo-Ibarra, Hugo Arasanz, Gonzalo Fernández-Hinojal, Gato-Cañas María).

## Abstract

The majority of lung cancer patients are refractory to PD-L1/PD-1 blockade monotherapy. This therapy may even accelerate progression and death in a group of patients called hyperprogressors. Here we demonstrate that the efficacy of PD-L1/PD-1 blockade therapy relies on baseline circulating highly-differentiated CD28^−^ CD27^−^ CD4 T cells (T_HD_ cells), which segregate patients in two non-overlapping groups. T_HD_ cells in cancer patients mostly comprised of central memory subsets that potently co-upregulated PD-1 and LAG3 upon antigen recognition. Low baseline T_HD_ numbers unequivocally identified intrinsic non-responders and hyperprogressors, whom aberrantly responded to therapy with a potent systemic proliferative T_HD_ cell burst. Responder patients showed significant reductions in systemic CD4 T_HD_ cells throughout therapy linked to expansion of the CD28+ CD27+ CD4 T cell compartment. Quantification of T_HD_ cells from peripheral blood samples prior to therapy allows identification of non-responders, hyperprogressors and responders, a critical issue in clinical oncology. These results place CD4 T cell responses at the center of anti-tumor immunity.

## Introduction

PD-L1/PD-1 blockade is demonstrating remarkable clinical outcomes since its first clinical application in human therapy (Brahmer et al., 2012; Topalian et al., 2012). These therapies interfere with immunosuppressive PD-L1/PD-1 interactions by systemic administration of blocking antibodies. PD-L1 is frequently overexpressed by tumor cells and correlates with progression and resistance to pro-apoptotic stimuli (Azuma et al., 2008; Gato-Canas et al., 2017; Juneja et al., 2017). PD-1 is expressed in antigen-experienced T cells and interferes with T cell activation when engaged with PD-L1 (Chemnitz et al., 2004; Karwacz et al., 2011).

A significant number of lung cancer patients are intrinsically refractory to PD-L1/PD-1 immune checkpoint blockade therapies. Indeed, there is accumulating evidence that PD-L1/PD-1 blockade might even have deleterious effects by accelerating disease and death in a group of patients called hyperprogressors (Champiat et al., 2017; Saada-Bouzid et al., 2017). Currently, there is no way of identifying them before the start of therapy. Hence, stratification of patients into hyperprogressors, non-responders and potential responders is of critical importance (Topalian et al., 2016) especially when the relevance of tumor PD-L1 expression as a predictive biomarker is still under debate, at least for some cancer types (Grigg and Rizvi, 2016). Other biomarkers such as the immunoscore (Galon et al., 2006), interferon gene signatures, mutational load and microsatellite instability require relatively large biopsies (Jamieson and Maker, 2017; Vranic, 2017). These techniques are often difficult to implement in a clinical context, and the retrieval of representative biopsies can sometimes be challenging.

In addition, the impact of T cell terminal differentiation in aged patients is often overlooked in cancer immunotherapies. Highly differentiated human T cells (T_HD_) accumulate with age by a progressive loss of CD27 and CD28 expression (Lanna et al., 2014). T cells are classified as poorly differentiated (CD28+ CD27+ T_PD_ cells), intermediately differentiated cells which lose CD27 expression (CD28+ CD27-T_INT_ cells) and T_HD_ cells (CD28-CD27-). In most cases T_HD_ cells correspond to terminally differentiated T cells with effector activities, also known as EMRA or senescent cells (Lanna et al., 2017; Lanna et al., 2014) (**Fig. 1A**). Although dysfunctional to some degree due to reduced expression of CD28 and CD27 co-receptors, these cells do constitute a large pool of antigen-specific T cells with potent effector activities when mobilized (Lanna et al., 2017; Lanna et al., 2014). Indeed, PD-1 blockade can recover some effector activities of CD8 T_HD_ cells *in vitro* (Henson et al., 2015). However, the clinical impact of PD-L1/PD-1 immune checkpoint inhibitor therapy in T_HD_ cells and its relationship with objective clinical responses have not been addressed yet. Here we focused on understanding the impact of T_HD_ cells over clinical responses in NSCLC patients undergoing PD-L1/PD-1 immune checkpoint blockade therapy.

## Results

### Baseline CD4 T_HD_ cells separate NSCLC patients in two non-overlapping groups

Systemic circulating T_HD_ cells were quantified in a cohort of 45 NSCLC patients and healthy age-matched donors (64.8 ± 8.3 *vs* 68.60 ± 8 years old, mean±standard deviation, SD). Cancer patients showed highly significantly increased numbers of CD4 T_HD_ cells (P<0.0001) but not of CD8 T_HD_ cells. Importantly, patients were distributed in two non-overlapping groups according to CD4 T_HD_ baseline numbers (**Fig. 1B**); Patients with above-average (G1 patients, 72.2% T_HD_ [67.2–76.8, 95% confidence interval (C.I.), N=21]) and below-average numbers (G2 patients, 28.2% T_HD_ [24–32.5, 95% C.I., N=24]). Accordingly, NSCLC patients showed very significantly lower numbers of CD28+ CD27+ CD4 T cells (P<0.001) (**Fig. 1B**). As CD4 T_HD_ cells segregated patients in two separate groups, we decided to characterize them at baseline and throughout treatment.

Surprisingly, CD4 T_PD_, T_INT_ and T_HD_ cells consisted of a mixture of differentiation phenotypes, including naïve/stem memory (CD62L+ CD45RA+), central memory (CM, CD62L+ CD45RA−), effector memory (EM, CD62L− CD45RA−) and effector cells (EF, CD62L^low/neg^ CD45RA+ or EMRA cells), independently on whether they came from G1 or G2 patients (**Fig. 1C, 1D**). CD4 T_HD_ were enriched in cells with central/effector memory phenotypes but not in EMRA cells, suggesting that these CD28^−^ CD27^−^ T cells were not truly terminally differentiated or senescent, although probably dysfunctional. To test this hypothesis, PD-1 expression was evaluated in systemic CD4 T_HD_ cells from NSCLC G1 and G2 patients. No significant PD-1 surface expression was observed in these cells (**Fig. 1E**). This was surprising as constitutive expression of markers such as PD-1 or LAG3 are trademarks of dysfunctional T cells in cancer. However, when CD4 T cells were stimulated by co-incubation with human lung A549 adenocarcinoma cells expressing a membrane-bound anti-CD3 antibody, CD4 T cells from NSCLC patients showed a strong co-up-regulation of PD-1 and LAG3 (**Fig. 1E**). A different expression pattern was observed without PD-1/LAG-3 co-expression when CD4 T cells from healthy donors were evaluated, in which PD-1 or LAG-3 represent activation markers. Moreover, PD-1 up-regulation was very significantly (P<0.001) pronounced in the more differentiated CD4 subsets, and especially in T cells from NSCLC patients (**Fig. 1F**).

**Figure 1.**
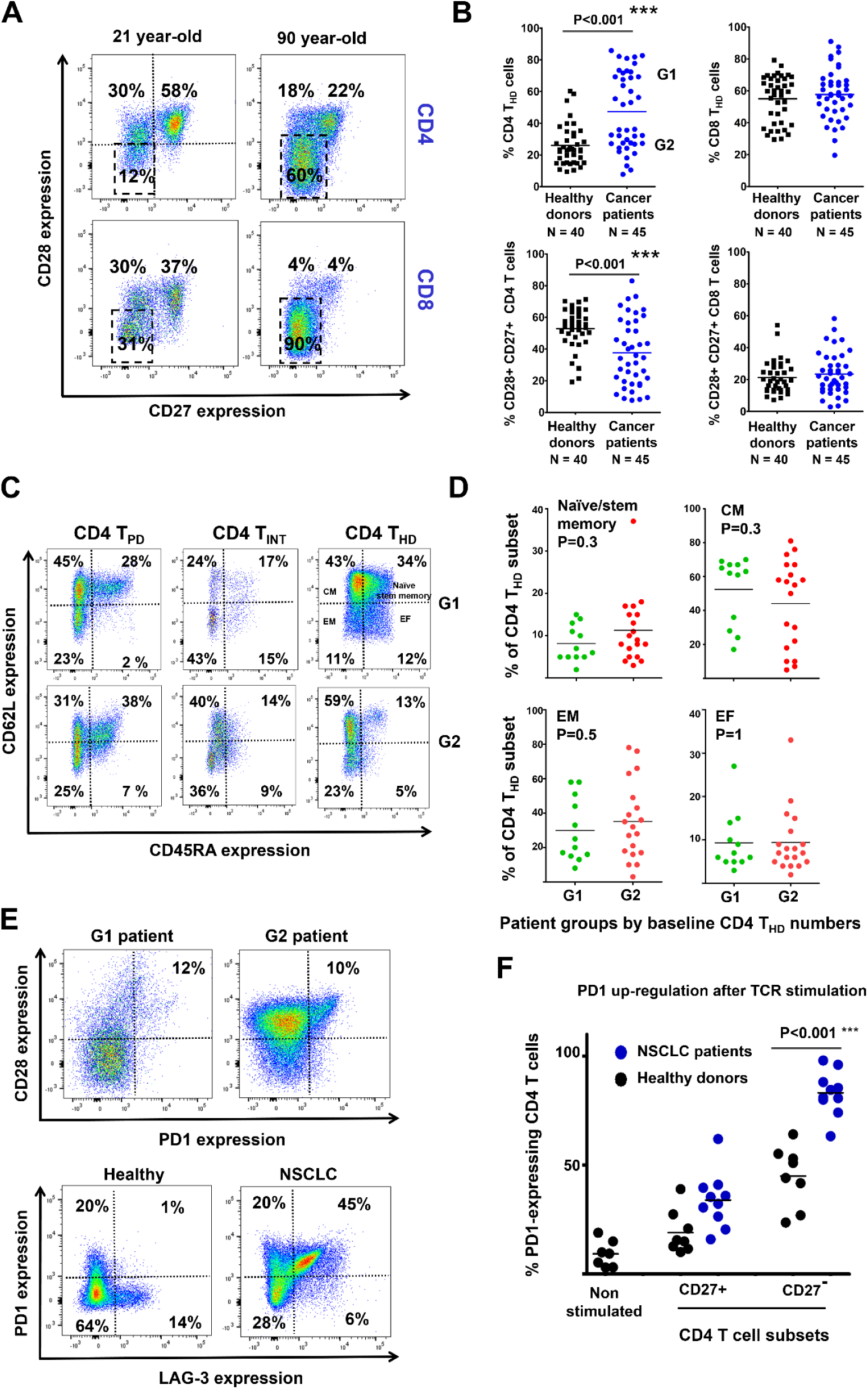
Systemic CD4 T_HD_ cells in NSCLC patients treated with PD-L1/PD-1 immune checkpoint inhibitors. **(A)** Flow cytometry density plots of CD4 (upper graphs) and CD8 T cells (lower graphs) from young (left graphs) and senior (right graphs) healthy donors, according to CD28-CD27 expression profiles. Dashed lines within the upper left graph show the gates used to quantify poorly differentiated (CD28+ CD27+), intermediately differentiated (CD28+ CD27−) and T_HD_ cells (CD28− CD27−). T_HD_ cells are highlighted in each graph by a square. Percentages of each cell subset are indicated within the graphs. **(B)** Circulating highly differentiated CD4 / CD8 (upper graphs), and poorly differentiated CD4 / CD8 (lower graphs) subsets in age-matched healthy donors or NSCLC patients before undergoing immunotherapies. G1 and G2, groups of patients classified according to high T_HD_ cells (G1) and low T_HD_ cells (G2). N, number of patients used for analyses. Relevant statistical comparisons are shown by the U of Mann-Whitney test. **(C)** Flow cytometry density graphs of CD4 T_HD_ from NSCLC G1 patients (upper graphs) or G2 patients (lower graphs) according to CD62L-CD45RA expression profiles. Dotted lines separate quadrants according to naïve/stem memory, central memory (CM), effector memory (EM) and effector phenotypes (EF), which include the percentage of cells in each quadrant. **(D)** As in (C) but representing data as scatter plot graphs for each patient classified according to G1 or G2 patient groups as indicated. Statistical comparisons performed by the U of Mann-Whitney. **(E)** Flow cytometry density plots of circulating CD4 T cells in G1 patients (upper left graph) and G2 patients (lower upper right graph) according to CD28-PD-1 expression profiles. The percentage of CD28+ PD-1+ CD4 T cells is indicated. The lower flow cytometry density graphs represent PD-1 and LAG3 up-regulation in CD4 T cells from a healthy donor (left graph) or an NSCLC patient (right graph) after T cell receptor (TCR) activation by A549 cells expressing a membrane bound anti-CD3 single-chain antibody. Percentages of cells within each quadrant are indicated. **(F)** Scatter plots representing the up-regulation of PD-1 after TCR activation as in (E) in healthy donors and NSCLC patients, separated into CD27+ and CD27-CD4 T cells. Relevant statistical comparisons are indicated, by the U of Mann Whitney. *** represents highly significant differences, respectively.

### Baseline CD4 T_HD_ numbers discriminate responses to PD-L1/PD-1 immune checkpoint blockade therapies

34 of the NSCLC patients continued with nivolumab, pembrolizumab or atezolizumab treatments following their current indications, and at the end of the study responders accounted to 20% (7 out of 34), consistent with the published efficacies for these agents (Herbst et al., 2016; Horn et al., 2017; Rittmeyer et al., 2017). To evaluate the impact of circulating CD4 T_HD_ over immunotherapies, we monitored CD4 T_HD_ cell numbers throughout therapy from routine small fresh blood samples. Strikingly, CD4 T_HD_ cell values before the start of therapy unequivocally discriminated patients according to responses (P=0.0008) (**Fig. 2A**). G2 patients (T_HD_ values below 40%) were all progressors [19 patients with 26.9% ± 7.8 baseline T_HD_ cells, (23–30.8, 95% C.I.)], while responders accounted to 47% of G1 patients with T_HD_ values above 40% [7 out of 15 patients with 71.5% ± 9 baseline T_HD_ cells, (63–80, 95% C.I.)]. Reciprocally, patients with CD28+ CD27+ CD4 T cell baseline values above 40% were all progressors (**Fig. 2B**) (P=0.005). Therefore, we defined patients with a “positive” baseline profile as those belonging to G1, while G2 represented patients with a “negative” baseline profile.

**Figure 2.**
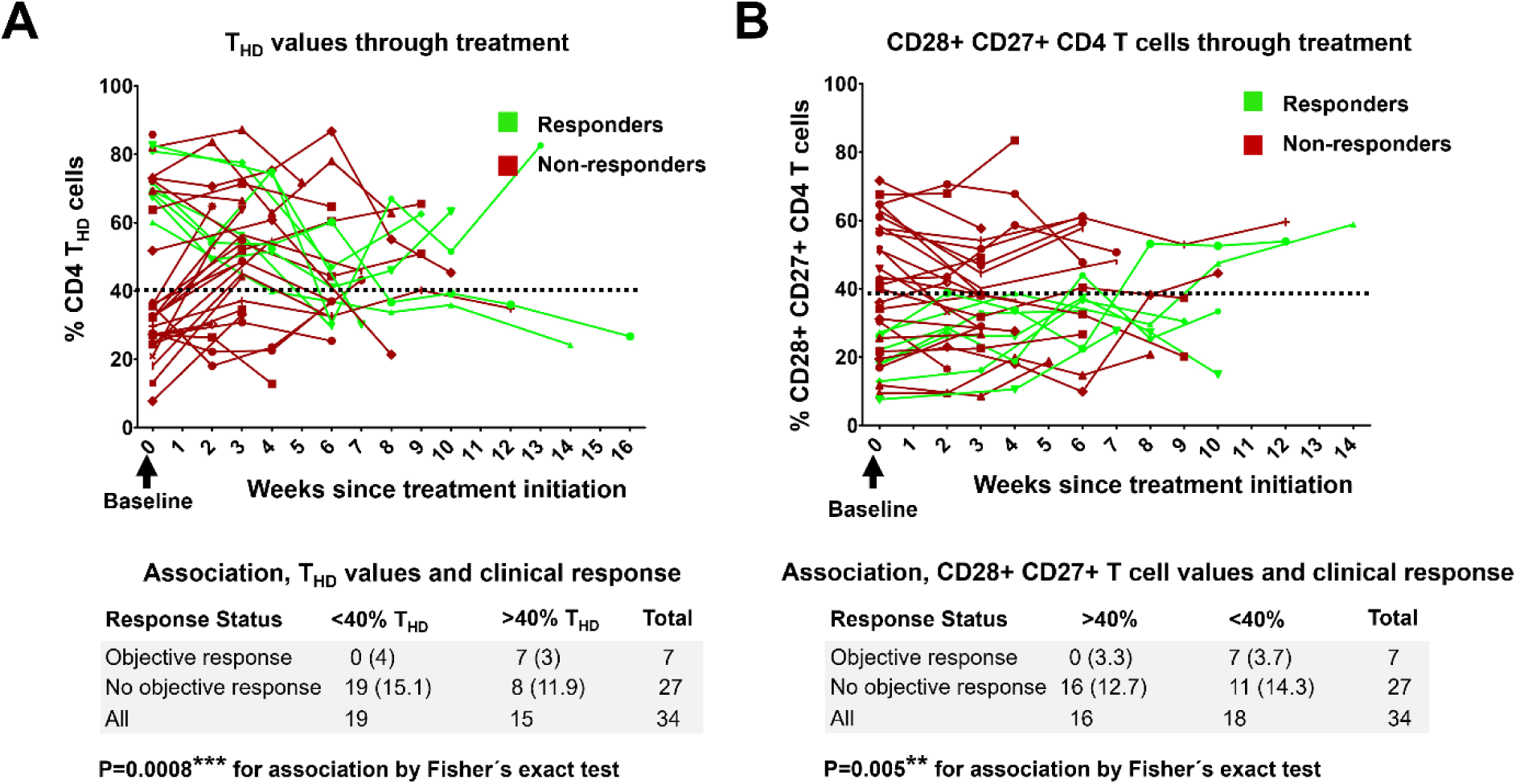
Dynamic changes of systemic CD4 T_HD_ and CD28+ CD27+ CD4 T cells throughout treatment. **(A)** Percentage of circulating CD4 T_HD_ cells in treated patients along therapy from baseline (arrow, time 0). In green, patients with objective responses. In red, non-responders. Dotted line, the lowest discriminating cut-off value (40%) separating G1 from G2 patients. No responders were observed below this cut-off value in the cohort study. Below the graph, correlation of responses to T_HD_ baseline values by the Fisher’s exact test. (B) Same as (A), but representing CD4 T_PD_ (CD28+ CD27+) cells.

### Immune checkpoint blockade therapy induces unique dynamic changes in circulating CD4 T_HD_ cells that correlate with clinical responses

Immune checkpoint inhibitors strongly affected CD4 T cell populations within the first cycle of treatment, and two main distinct dynamic profiles were identified. Pattern 1 or “T_HD_ burst” consisted in a highly significant increase in systemic CD4 T_HD_ cells [12.4% increase, (6.2, 18.5) 95% CI, N=27, P<0.0001, one-tailed paired t test)] and was associated to tumor progression without exception in our cohort of patients (**Fig. 3A, 3C, 3D**). Pattern 2 or T_HD_ decrease was characterized by very significant reduction in systemic T_HD_ cells [−14.4%, (−8, −21), 95% CI, N=7, P<0.0001], concomitant to an expansion of CD28+ CD27+ CD4 T cells and primarily associated to tumor regression (**Fig. 3B, 3C, 3D**). There was a very highly significant correlation (P=0.0001) between T_HD_ changes and therapeutic outcome (**Fig. 3D**).

**Figure 3.**
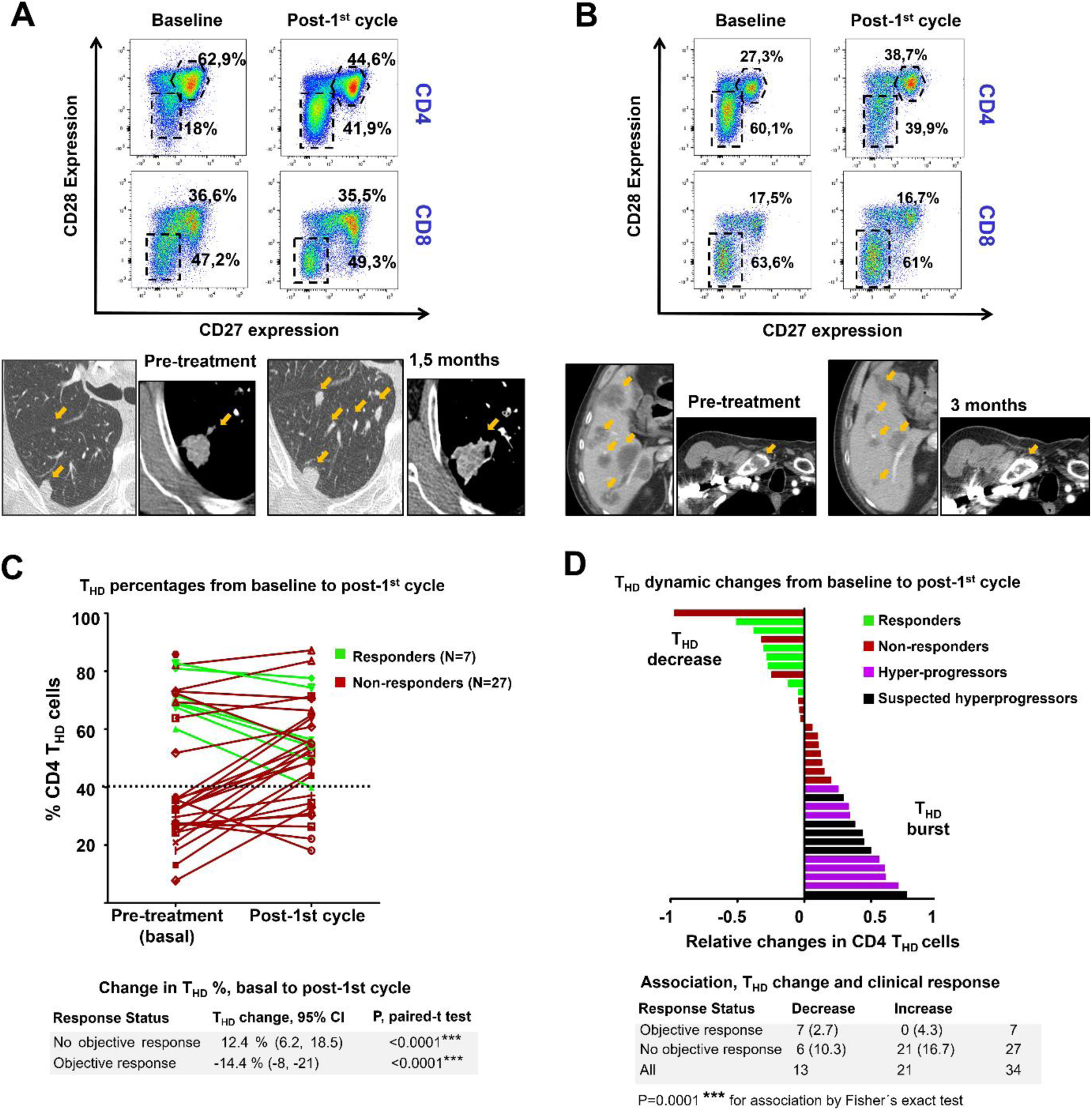
Systemic CD4 T_HD_ dynamic changes and clinical responses. **(A)** Clinical case of a progressor with a G2 baseline profile associated to a “T_HD_ burst” response. Flow cytometry density plots show baseline and post-first cycle CD4 (upper graphs) and CD8 T cells (lower graphs). Highly differentiated and poorly differentiated CD4 T cells are highlighted within doted square gates and hexagonal gates, respectively, together with their percentage. Below, CT scans of lung metastases and primary tumor progressing from baseline (left scans) after one month and a half (right scans) of therapy. Lesions are indicated with arrows. **(B)** Clinical case of a responder with a G1 baseline profile associated to systemic T_HD_ reduction. CT scans show regression of hepatic and clavicular bone metastases (indicated with arrows) before (left scans) and after 3 months (right scans) of therapy. **(C)** Change in circulating CD4 T_HD_ cells from baseline (pre-treatment) to post-first cycle of therapy. In red, progressors, in green objective responders. The 40% cut-off value separating G1 from G2 patients is shown. Below, increase and decrease in T_HD_ cells within reponders or non-responders, as tested by paired t tests. **(D)** Waterfall plot of the relative changes in CD4 T_HD_ cells in each patient from baseline to post-first cycle of therapy. Green bars, patients with objective responses; red bars, non-responders; purple, radiologically-confirmed hyperprogressors; black, suspected hyperprogressors. Below, correlation of T_HD_ cell change with clinical outcome by the Fisher’s exact test.

To find out if T_HD_ bursts were the result of active proliferation of CD4 T_HD_ cells from baseline to post-first cycle of therapy, Ki67 expression was analyzed by intracellular flow cytometry in T_HD_ cells from non-responder and responder patients **(Fig. 4A, 4B)**. T_HD_ cells in non-responders showed a significant increase in Ki67 expression. In contrast, responders showed a markedly decrease in T_HD_ Ki67 expression from baseline to post-first cycle of therapy, together with increased Ki67 expression in the CD28+ CD27+ CD4 T cell compartment **(Fig. 4C)**. These results indicated that T_HD_ bursts were likely the result of systemic T_HD_ proliferation.

**Figure 4.**
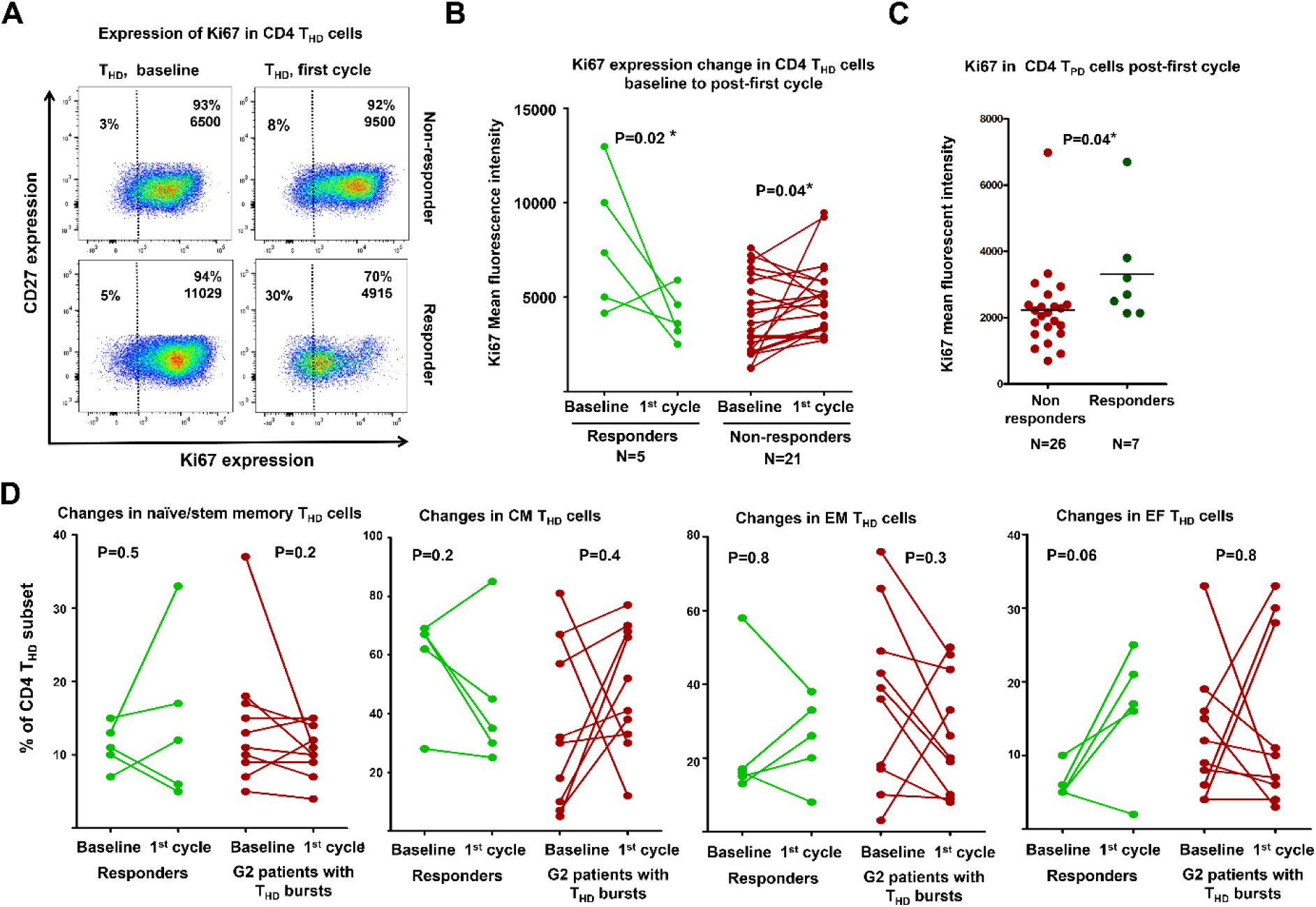
Proliferation of CD4 T_HD_ in responders and non-responders. **(A)** Flow cytometry density plots of Ki67 expression in T_HD_ cells from a progressor (upper two graphs) and a responder (lower two graphs) at baseline and post-first cycle of therapy as indicated. Percentage and Ki67 mean fluorescent intensities in proliferating T_HD_ cells are indicated within the graphs. **(B)** Change in Ki67 expression in CD4 T_HD_ cells from responders and non-responders, as indicated. Only data was plotted from patients in which baseline and first-cycle Ki67 values were available. Paired t-tests were performed to compare the change from baseline to post-first cycle of therapy. **(C)** Dot plots of Ki67 expression in CD28+ CD27+ CD4 T cells (right graph) in non-responders and responders as indicated, in our cohort study. Differences were tested by the U of Mann-Whitney test. **(D)** Dot plots of changes in the percentage of circulating T_HD_ differentiation subsets (as indicated) from baseline to post-first cycle of therapy, in patients exhibiting T_HD_ bursts compared to responders. Data from patients with available CD62L-CD45RA profiles were used in the analyses. Relevant statistical comparisons are shown within the graphs, using paired t tests; N, number of patients used in the analyses; * indicate significant differences.

To test whether T_HD_ bursts occurred in specific subsets, the changes from baseline to post-first cycle of therapy were studied only in patients exhibiting T_HD_ bursts, and compared to those from responder patients. Although the changes were not significant either in responders (with T_HD_ decrease) or non-responders exhibiting T_HD_ bursts with the current number of patients, there were clear trends. CD4 T_HD_ bursts were enriched in CM subsets in detriment of more differentiated effector subsets (EM and EF cells) **(Fig. 4D)**. In contrast, CD4 T_HD_ cells that remained in responders post-first cycle of therapy were enriched in EM and EF cells.

### CD4 T_HD_ bursts define primary resistance to PD-L1/PD-1 blockade and hyperprogression

All G2 patients showed tumor progression **(Fig. 5A)**. Within this group, hyperprogressors were identified following the definition by Sâada-Bouzid *et al* (Saada-Bouzid et al., 2017) but using as a threshold a tumor growth kinetics ratio equal or superior to 5 **(Fig. 5B)**. We confirmed that a negative T_HD_ baseline profile significantly correlated with radiologically-confirmed hyperprogressors (P=0.01) **(Fig. 5C)** whom showed highly significant T_HD_ bursts (P=0.0001) following the first cycle of therapy **(Fig. 5D)**. Six patients were identified as probable hyperprogressors by clinical parameters, whom either died before radiological confirmation or the disease was not evaluable by radiological criteria. Their immunological profiles were consistent with radiologically-confirmed hyperprogressors **(Fig. 5C and 5D)**. All of them experienced early progression of disease compared to the rest (median progression-free survival (mPFS)=6 weeks [5.7–6.3, 95% C.I.] *versus* 8.9 weeks [4.6–13.1, 95% C.I.], p=0.002).

The agreement between the radiological criterium and the immunological profiling was significant in the identification of hyperprogression by a kappa test (κ=0.742). Hence, a G2 profile associated to significant “T_HD_ bursts” objectively characterized hyperprogressive disease in NSCLC patients without being influenced by previous tumor burden or dynamics.

**Figure 5.**
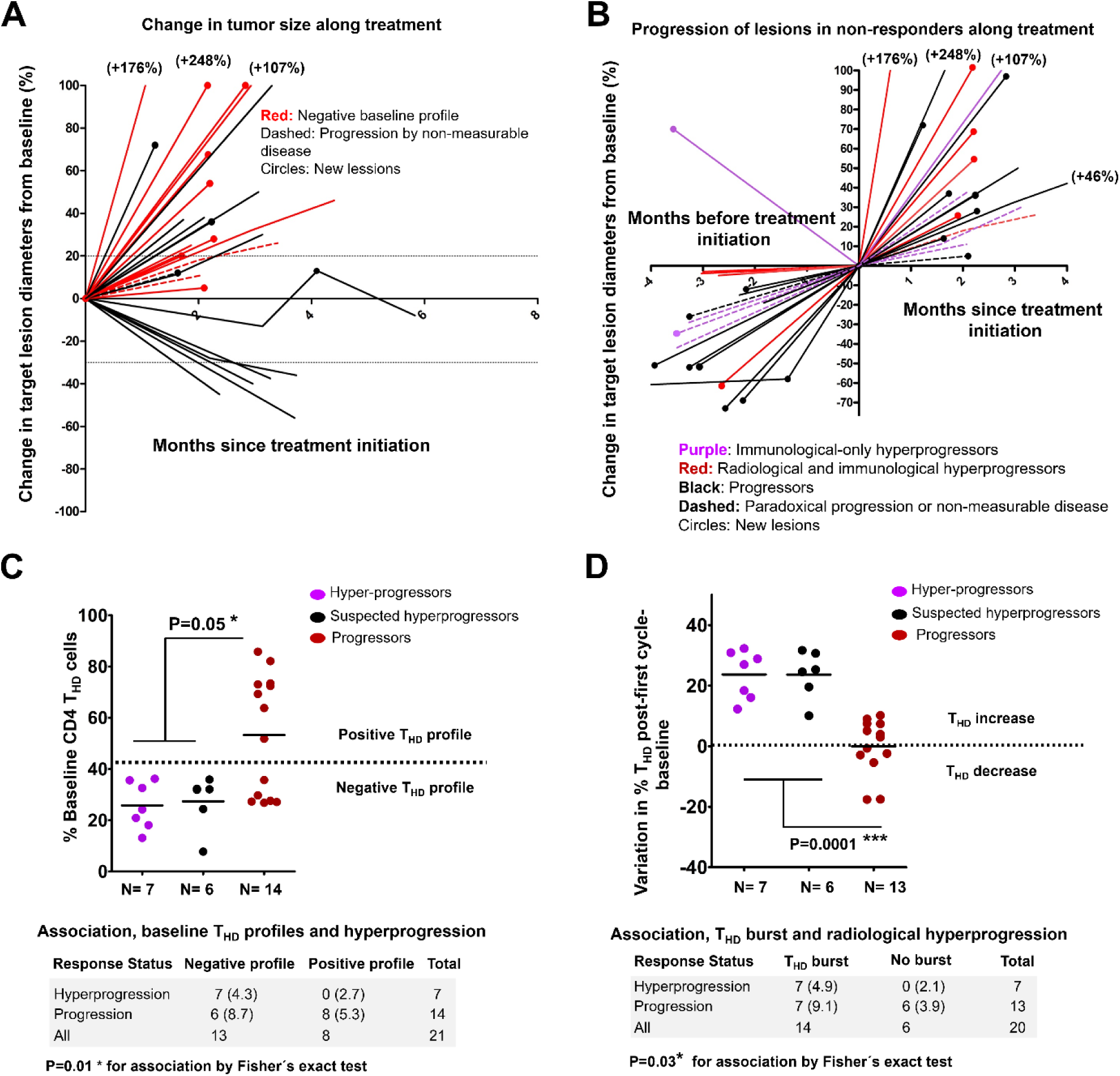
T_HD_ immunological profiles and hyperprogressive disease. **(A)** Spider plot of change in target lesions. Red, patients that started therapy with a negative baseline profile. These patients presented progressive disease, or growth of lesions. **(B)** Spider plot of change in target lesions of progressors before and after the start of immunotherapy. **(C)** Scatter plot of baseline T_HD_ cell values in hyperprogressors, suspected hyperprogressors and progressors, as indicated. Dotted line shows the 40% cut-off value separating G1 from G2 patients. Below, correlation of baseline T_HD_ cells with radiologically-confirmed hyperprogressors by a Fisher’s exact test. Suspected hyperprogressors were excluded. **(D)** Scatter plot of changes in CD4 T_HD_ percentage from baseline to post-first cycle of therapy in radiologically-confirmed hyperprogressors, suspected hyperprogressors and progressors. Dotted line separates T_HD_ increases from decreases. Differences were tested by U of Mann-Whitney test. Suspected hyperprogressors were excluded. Below, correlation of T_HD_ burst with radiologically-confirmed hyperprogressors by a Fisher’s exact test. N, number of patients in each group; Comparisons of CD4 T_HD_ cells and changes in CD4 T_HD_ cells were performed with the U of Mann-Whitney excluding suspected hyperprogressors; N, number of patients used in the analyses; *,**, ** indicates significant, very and highly significant differences.

### Baseline T_HD_ numbers constitute a strong and reliable predictive biomarker for primary resistance and hyperprogression with practical clinical application

To improve specificity even at the cost of a lower sensitivity for identification of intrinsic non-responders, the negative baseline T cell profile was refined as the combination of CD4 T_HD_ <40% and CD28+ CD27+ CD4 T cells > 40%. In patients with this strict negative baseline profile, the mPFS was only 6.0 weeks (95% C.I., 5.7 to 6.3) **(Fig. 6A)**. A comparison of negative and non-negative baseline profiles showed hazard ratios for disease progression or death that favored the latter [3.39 (1.4–8.17; 95% C.I.) P = 0.007] **(Fig. 6A)**. Patients were also stratified into hyperprogressors according to the immunological criteria defined in this study, with an mPFS of only 6 weeks (5.7–6.3; 95% C.I.). Hazard ratios for disease progression or death favored patients with non-hyperprogressive T cell dynamics [3.89 (1.58–9.58; 95% C.I.) P = 0.002] **(Fig. 6B)**. To discard that the baseline T_HD_ profile was a prognostic instead of a predictive factor, the median time elapsed from diagnosis to enrolment for immunotherapy was evaluated in patients with negative or positive T_HD_ profiles. There were no significant differences between the two groups, confirming the predictive and not the prognostic value of T_HD_ profiling (10.9 months vs 9.8 months, P=0.9899) **(Fig. 6C)**. ROC analysis was performed in our patient cohort to test the robustness of T_HD_ quantification prior to therapy as a predictive biomarker **(Fig. 6D)**. The association of baseline T_HD_ cells with clinical output was very significant (R=0.84, P=0.006), with a cut-off value of <57.7% to achieve 100% specificity and 75% sensitivity.

**Figure 6.**
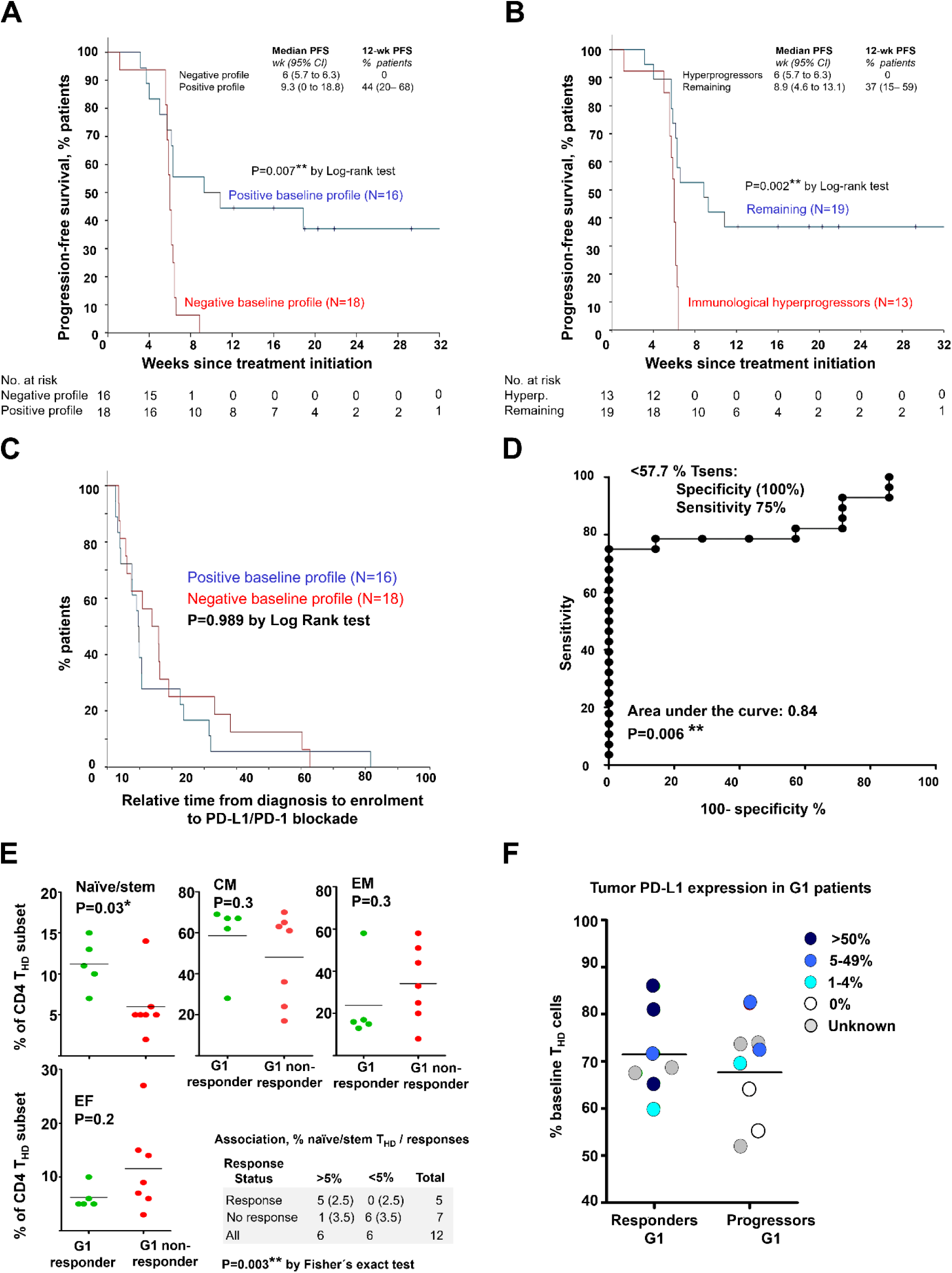
T_HD_ profiling as a predictive biomarker of responses. **(A)** Kaplan-Meier plot for PFS in patients undergoing immune checkpoint inhibitor therapies stratified by strict baseline negative and positive T cell profiles as defined in the text. Patients starting therapy with a negative baseline profile had an overall response rate (ORR) of 0% and all experienced progression or death by week 9. ORR was 38.9% for patients with a positive baseline profile, and the 12-week PFS was 44%. **(B)** As in (A) but stratifying patients according to hyperprogression assessed by immunological criteria (negative CD4 T_HD_ baseline profile-significant T_HD_ burst). Patients classified as immunological progressors progressed or died by week 7. **(C)** Kaplan-Meier plot of time of diagnosis to enrolment in patients stratified by positive or negative CD4 T_HD_ profiles as indicated, demonstrating no significant prognostic value. **(D)** ROC analysis of baseline CD4 T_HD_ quantification as a predictive biomarker. Within the graph, highest cut-off value of CD4 T_HD_ cells to discriminate intrinsic responders with 100% specificity. **, indicates very significant differences. **(E)** Scatter plots of percentages of baseline T_HD_ differentiation subsets as indicated on top of each graph in responders and non-responders from G1 patients. Statistical comparisons were performed with the U of Mann-Whitney test. Right bottom, correlation of the percentage of naïve/stem memory CD4 T_HD_ cells with objective responses in G1 patients by a Fisher’s exact test. **(F)** Scatter plot of the percentage of baseline CD4 T_HD_ cells according to tumor expression levels as shown in the legend.

### Identification of responders by baseline T_HD_ subset profiling and PD-L1 tumor status

Identification of responders with a high probability prior to therapy is currently a major challenge. In this study, all responders belonged to G1 patients and presented a specific T_HD_ fingerprint consisting of higher numbers of naïve/stem memory CD4 T_HD_ cells compared to G1 non-responders (P=0.03) **(Fig. 6E)**. There was a very significant association between G1 patients with naïve/stem memory T_HD_ cells above 5% and objective responses (P=0.003).

PD-L1 tumor expression levels could be evaluated in 21 of the patients before therapy, and did not significantly correlate with baseline T_HD_ G1 or G2 profiles (P=0.1). PD-L1 expression correlated with objective responses at the limit of statistical significance in our cohort study when used as a single stratifying factor (P=0.052 by Fishers exact test) (data not shown). PD-L1 tumor expression seemed to segregate G1 patients into responders and non-responders **(Fig. 6F)**, in our limited cohort of patients in whom PD-L1 tumor expression levels could be evaluated. There was a tendency for G1 responder patients compared to G1 non-responders to have higher PD-L1 tumor expression prior to therapy. These results suggested that patients with baseline naïve/stem memory T_HD_ cells above 5% together with PD-L1 tumor positivity may accurately identify responders amongst the G1 patient population.

## Discussion

Our data shows that the efficacy of PD-L1/PD-1 blockade monotherapies in metastasic NSCLC patients heavily relies on systemic CD4 T_HD_ cell numbers. Importantly, we are unequivocally identifying patients in our clinical practice with primary resistance and a high risk of hyperprogressive disease before enrolment, by quantification of T_HD_ cells from routine small blood samples. While there has been a recent output of potential biomarkers from blood sampling, most of them have prognostic value rather than predictive capacities, while others are rather challenging to implement in routine clinical practice.

CD8 T cell subsets were extensively studied with similar results as reported (Kamphorst et al., 2017) but without practical stratification capacities. The CD8 T cell response was delayed and followed CD4 T_HD_ dynamic changes but at a lesser extent. Therefore, CD8 T cell monitoring had a lack of practical predictive capacities. Initially, we hypothetized that CD4 T_HD_ cells in cancer patients were senescent, terminally-differentiated T cells, a subset strongly associated to the EMRA phenotype (Lanna et al., 2017; Lanna et al., 2014). To our surprise, CD4 T_HD_ cells in cancer patients were enriched in central and effector memory subtypes, strongly suggesting that T_HD_ cells did not reach terminal differentiation. Indeed, CD4 T_HD_ cells expressed high levels of Ki67, suggesting that they were proliferating in cancer patients. This raises the question of whether reduced CD28 and CD27 expression in T cells is a true hallmark of T cell senescence, at least in lung cancer patients. Nevertheless, the more differentiated CD4 T cell subsets in lung cancer patients strongly co-expressed PD-1 and LAG3 after stimulation compared to CD28+ CD27+ T cells. This suggests that the systemic pool of differentiated CD4 T cells is dysfunctional in patients, and prone to inactivation by PD-L1/PD-1 interactions. NSCLC patients were separated in two distinct groups according to baseline CD4 T_HD_ numbers, strongly suggesting that lung cancer patients had diametrically different immunological responses to cancer. Responder patients presented significantly higher CD4 T_HD_ baseline numbers which may represent a pre-existing large pool of antigen-specific central and effector memory T cells with potential anti-tumor capacities. Responders showed decreases in circulating CD4 T_HD_ cells following antibody administration, which may indicate either a recovery of CD28-CD27 expression, or mobilization from peripheral blood to secondary lymphoid organs/tumor sites. In contrast, a systemic expansion of central memory T_HD_ cells (most likely through active proliferation) was always associated to non-responders which was plainly apparent in hyperprogressors. A negative baseline profile associated to “T_HD_ bursts” accurately correlate with hyperprogressive disease in NSCLC without the need of obtaining radiological evidence, which supports hyperprogression as a true phenomenon with a biological basis.

According to our results in our cohort study, NSCLC patients with a negative baseline profile (G2 patients) do not respond to PD-L1/PD-1 immune checkpoint blockade monotherapy. Their identification prior to therapy helps avoiding the enrolment of hyperprogressors and intrinsic non-responders. Of the remaining patients, quantification of CD4 T_HD_ cells with naïve/stem memory phenotypes coupled to tumor expression of PD-L1 can help identifying responders with accuracy. Nevertheless, administration of therapy would not be indicated if T_HD_ bursts are observed following the first cycle of therapy.

T_HD_ profiling can be performed in patients some time before enrolment into immunotherapies without necessarily quantifying this subset right before starting therapy. Indeed, our immunological profiling helps to make objective decisions regarding NSCLC patients under PD-L1/PD-1 blockade, and places CD4 T cell responses at the center of anti-tumor immunity.

## Materials and Methods

### Study Design

The study was approved by the Ethics Committee at the Hospital Complex of Navarre, and strictly followed the principles of the Declaration of Helsinki and Good Clinical Practice guidelines. Written informed consent was obtained from each participant, and samples were collected by the Blood and Tissue Bank of Navarre, Health Department of Navarre, Spain. 27 patients diagnosed with non-squamous and 7 with squamous NSCLC were recruited at the Hospital Complex of Navarre (**Table S1 in supplementary material**). Patients had all progressed to first line chemotherapy or concurrent chemo-radiotherapy. Eligible patients were 18 years of age or older who agreed to receive immunotherapy targeting PD-1/PD-L1 following the current indications **(table S1)**. Tumor PD-L1 expression could be quantified in 21 of these patients before the start of therapies. Measurable disease was not required. The exclusion criteria consisted on concomitant administration of chemotherapy or previous immunotherapy treatment. NSCLC patients had an age of 64.8 ± 8.3 (mean ± standard deviation, S.D., N=45). Age-matched healthy donors were recruited from whom written informed consent was also obtained, with an age of 68.60 ± 8 (mean ± S.D., N=40).

Therapy with nivolumab, pembrolizumab and atezolizumab was provided following current indications (Herbst et al., 2016; Horn et al., 2017; Rittmeyer et al., 2017). 4 ml peripheral blood samples were obtained prior and during immunotherapy before administration of each cycle. PBMCs were isolated as described (Escors et al., 2008) and T cells analysed by flow cytometry. The participation of each patient concluded when a radiological test confirmed response or progression, with the withdrawal of consent or after death of the patient. Tumor responses were evaluated according to RECIST 1.1 (Eisenhauer et al., 2009) and Immune-Related Response Criteria (Wolchok et al., 2009). Hyperprogression was identified by the criteria proposed by Saada-Bouzid *et al*. (Champiat et al., 2017; Saada-Bouzid et al., 2017). Objective responses were confirmed by at least one sequential tumor assessment.

### Flow cytometry

Surface and intracellular flow cytometry analyses were performed as described (Gato-Canas et al., 2017; Karwacz et al., 2011). T cells were immediately isolated and stained. 4 ml blood samples were collected from each patient, and PBMCs isolated by FICOL gradients right after the blood extraction. PBMCs were washed and cells immediately stained with the indicated antibodies in a final volume of 50 μl for 10 min in ice. Cells were washed twice, resuspended in 100 μl of PBS and analyzed immediately. The following fluorochrome-conjugated antibodies were used at the indicated dilutions: CD4-FITC (clone M-T466, reference 130–080–501, Miltenyi Biotec), CD4-APC-Vio770 (clone M-T466, reference 130–100–455, Miltenyi Biotec), CD4-PECy7 (clone SK3, reference 4129769, BD Biosciences,) CD27-APC (clone M-T271, reference 130–097–922, Miltenyi Biotec), CD27-PE (clone M-T271, reference 130–093–185, Miltenyi Biotec), CD45RA-FITC (reference 130–098–183, Miltenyi Biotec), CD62L-APC (reference 130–099–252, Miltenyi Biotech), CD28-PECy7 (clone CD28.2, reference 302926, Biolegend), PD-1-PE (clone EH12.2H7, reference 339905, Biolegend), CD8-FITC (clone SDK1, reference 344703, Biolegend), CD8-APC-Cy7(clone RFT-8, reference A15448, Molecular probes by Life technologies).

### Cell culture

Human lung adenocarcinoma A549 cells were a kind gift of Dr Ruben Pio, and were grown in standard conditions. These cells were modified with a lentivector encoding a single-chain version of a membrane bound anti-OKT3 antibody.

### Spanning-tree progression analysis of density-normalized events (SPADE)

Flow cytometry data was analyzed with SPADE v3 (Qiu et al., 2011) using forward, scatter, CD4, CD28 and CD27 expression as overlapping markers for cell clustering. Arcsinh with cofactor of 150 was used for data transformation, from a maximum of 100000 events and 30 clusters.

### Data collection and Statistics

T cell percentages were quantified using Flowjo (Lanna et al., 2017; Lanna et al., 2014). The percentage of CD4/CD8 T_HD_ (CD28 and CD27 double-negative) and poorly differentiated T cells (CD28+ CD27+) were quantified prior to therapy (baseline), and before administration of each cycle of therapy. Gates in flow cytometry density plots were established taking non-senescent T cells as a reference. Data was recorded by M.Z.I., and separately analyzed thrice by M.Z.I. and H.A.E. independently. Cohen’s kappa coefficient was utilized to test the inter-rater agreement in classification of immunological profiles (κ=0.939).

The mode of action, pharmacokinetics, adverse events and efficacies of the three PD-L1/PD-1 blocking agents are comparable in NSCLC, which act through the interference with the inhibitory interaction between PD-L1 and PD-1 (Herbst et al., 2016; Horn et al., 2017; Rittmeyer et al., 2017). Treatments administered to the patients were allocated strictly on the basis of their current indications, and independently of any variable under study. Hence, in the study design the use of all data was pre-specified to be pooled to enhance statistical power, and thereby reducing type I errors from testing the hypotheses after *had hoc* subgrouping into specific PD-L1/PD-1 blockers. The number of patients assured statistical power for Fisher’s exact test of 0.95 and superior for Student t and Mann-Whitney tests (G*Power calculator) (Faul et al., 2009), taking into account that the expected proportion of responders using any of the three PD-L1/PD-1 immune checkpoint blockade drugs in NSCLC is around 20% to 25% (if no stratification using PD-L1 tumor expression levels is used) (Herbst et al., 2016; Horn et al., 2017; Rittmeyer et al., 2017). Our study confirmed the therapeutic efficacy of these agents by achieving a 20% response rate. Two pre-specified subgroup analyses in the study were contemplated. The first, baseline T cell values; the second, post-first cycle T cell changes from baseline. The study protocol contemplated the correlation of these values with responses using Fisher’s exact test, paired Student t tests/repeated measures ANOVA (if normally distributed) or U of Mann-Whitney/Kruskal-Wallis (if not normally distributed, or data with intrinsic high variability) to be correlated with responses. Two-tailed tests were applied with the indicated exceptions (see below). Importantly, the treatment administered to the patients was allocated independently of their basal immunological profiles, and strictly followed the current indications for the PD-L1/PD-1 inhibitors.

The percentage of T cell subsets in untreated cancer patients was normally distributed (Kolmogorov-Smirnov normality test), but not in age-matched healthy donors. Hence, to compare T cell values between two independent cancer patient groups, two-tailed unpaired Student t tests were used while comparisons between healthy subjects and cancer patients were carried out with the U of Mann-Whitney. The mean age of cancer patients and healthy donors was 64.8 ± 8.3 (S.D.) and 68.60 ± 8 (S.D.), respectively. Percentages of T cell populations in treated patients were not normally distributed, so response groups were compared with either Mann-Whitney (comparisons between two independent groups) or Kruskall-Wallis for multi-comparison tests if required and as indicated in the manuscript. To confirm the increase or decrease in T_HD_ cells between baseline and post-first cycle of therapy in either responders or non-responders, one-tailed paired t tests were carried out. To test changes in Ki67 expression in T cells between baseline and post-first cycle of therapy in either responders or non-responders, two-tailed paired t tests were carried out. Fisher’s exact test was used to assess the association between CD4 T_HD_ dynamic profiles or the baseline values of T_HD_ cells with clinical responses. *Post hoc* Cohen’s kappa coefficient test was used to test the agreement of radiological versus immunological criteria for the identification of hyperprogressors.

Progression free survival (PFS) was defined as the time from the starting date of therapy to the date of disease progression or the date of death by any cause, whichever occurred first. PFS was censored on the date of the last tumor assessment demonstrating absence of progressive disease in progression-free and alive patients. PFS rates at 12 and 28-weeks was estimated as the proportion of patients who were free-of-disease progression and alive at 12 and 28 weeks after the initiation of immunotherapies. Patients who dropped out for worsening of disease and did not have a 28-week tumor assessment were considered as having progressive disease. Overall response rate (ORR) was the proportion of patients who achieved best overall response of complete or partial responses.

PFS was represented by Kaplan-Meier plots and long-rank tests utilized to compare cohorts. Hazard ratios were estimated by Cox regression models. For PFS analyses, patients were pre-specified to be stratified only on the basis of their basal T_HD_ values to avoid increase in type I errors due to multiplicity by subgroup analyses. Receiver operating characteristic (ROC) analysis was performed with baseline T_HD_ numbers and response/non-response as a binary output. Statistical tests were performed with GraphPad Prism 5 and SPSS statistical packages.

## Acknowledgments

We sincerely thank the patients and families that generously agreed to take part in this study. We are thankful to Drs Luis Montuenga and Ruben Pio for their constructive comments and input.

## Funding

This research was supported by Asociación Española Contra el Cáncer (AECC, PROYE16001ESCO); Instituto de Salud Carlos III, Spain (FIS project grant PI17/02119), a “Precipita” Crowdfunding grant (FECYT). D.E. is funded by a Miguel Servet Fellowship (ISC III, CP12/03114, Spain); M.Z.I. is supported by a scholarship from Universidad Pública de Navarra; H.A. is supported by a scholarship from AECC; M.G.C. is supported by a scholarship from the Government of Navarre.

## Author contributions

M.Z.I. designed and carried out experiments, collected data, analyzed data. H. A.E. designed and carried out experiments, collected data, analyzed data. G.F.H. recruited patients, collected data, analyzed clinical data. M.G.C. carried out experiments, collected data, analyzed data. B.H.M. recruited patients, collected data, analyzed clinical data. M.M.A. recruited patients, collected data, analyzed clinical data. M.J.L. recruited patients, collected data, analyzed clinical data. A.F. recruited patients, collected data, analyzed clinical data. L.T. recruited patients, collected data, analyzed clinical data. R.V. supervised the clinical staff, recruited patients, analyzed clinical data. G.K. conceived the project, supervised non-clinical researchers, analysed data and wrote the paper. D.E. conceived the project, supervised non-clinical researchers, analysed data and wrote the paper. All authors participated in the writing of the manuscript.

## Competing interests

The authors declare no competing interests.

## Supplementary Materials

**Table S1.** Baseline patient characteristics

| **CHARACTERISTIC** | **Non-Small Cell Lung Cancer (N = 34)** |
| --- | --- |
| **Age category – no. (%)** |  |
| 41-50 | 1 (2.9%) |
| 51-60 | 10 (29.4%) |
| 61-70 | 15 (44.1%) |
| 71-80 | 7 (20.6%) |
| >80 | 1 (2.9%) |
| **Sex – no. (%)** |  |
| Male | 25 (73.5%) |
| Female | 9 (26.5%) |
| **ECOG Performance-status score – no. (%)** |  |
| 0-1 | 26 (76.5%) |
| 2-4 | 8 (23.5%) |
| **Smoking habit – no. (%)** |  |
| Yes | 31 (91.2%) |
| No | 3 (8.8%) |
| **Metastases location – no. (%)** |  |
| Lymph nodes | 3 (8.8%) |
| Viscera | 3 (8.8%) |
| Both | 28 (82.4%) |
| **Immunotherapy – no. (%)** |  |
| Pembrolizumab | 4 (11.8%) |
| Nivolumab | 13 (38.2%) |
| Atezolizumab | 17 (50%) |
| **Treatments received during 3 months prior to immunotherapy – no. (%)** |  |
| Platinum-based chemotherapy | 11 (32.3%) |
| Other chemotherapy | 12 (35.3%) |
| Concurrent chemoradiotherapy | 1 (2.9%) |
| Radiotherapy | 2 (5.9%) |
| Targeted therapies | 0 (0%) |
| None | 8 (23.5%) |
| **Lymphocyte count – no. (%)** |  |
| < 1.5 x 10<sup>9</sup>/L | 8 (23.5%) |
| 1.5 – 4.0 x 10<sup>9</sup>/L | 26 (76.5%) |
| **Tumor PD-L1 expression - no. (%)** |  |
| 0% | 9 (26.5%) |
| 1-4% | 4 (12.8%) |
| 5-49% | 4 (12.8%) |
| ≥ 50% | 5 (14.7%) |
| Non-evaluable | 12 (35.3%) |
| **Baseline lymphocyte profile** |  |
| Disfavorable | 16 (47%) |
| Non-disfavorable | 18 (53.5%) |

